# Predicting input signals of transcription factors in *Escherichia coli* through metabolomics and transcriptomics

**DOI:** 10.1101/2025.02.24.639830

**Authors:** Julian Trouillon, Alexandra E. Huber, Yannik Trabesinger, Uwe Sauer

## Abstract

The activity of bacterial transcription factors (TFs) is typically modulated through direct interactions with small molecules. However, these input signals remain unknown for most TFs, even in well-studied model bacteria. Identifying these signals typically requires tedious experiments for each TF. Here, we develop a systematic workflow for the identification of TF input signals in bacteria based on metabolomics and transcriptomics data. We inferred the activity of 173 TFs from published transcriptomics data and determined the abundance of 279 metabolites across 40 matched experimental conditions in *Escherichia coli*. By correlating TF activities with metabolites abundances, we successfully identified previously known TF-metabolite interactions and predicted novel TF effector metabolites for 41 TFs. To validate our predictions, we conducted *in vitro* assays and confirmed a predicted effector metabolite for LeuO. As a result, we established a network of 80 regulatory interactions between 71 metabolites and 41 *E. coli* TFs. This network includes 76 novel interactions that encompass a diverse range of chemical classes and regulatory patterns, bringing us closer to a comprehensive TF regulatory network in *E. coli*.

## Introduction

Adapting to changing environmental conditions requires adjustments in the cellular proteome. In bacteria, these adjustments are primarily mediated by transcription factors (TFs), which regulate gene expression through their interactions with DNA (Browning & Busby, 2016; Mejía-Almonte *et al*, 2020; Fang *et al*, 2017; Panis *et al*, 2015). To elicit appropriate gene expression responses, information from the extracellular environment or the cellular state must be relayed to TFs. The predominant mechanism for this information transfer in bacteria is the allosteric binding of intracellular metabolites (Ledezma-Tejeida *et al*, 2021; Yugi & Kuroda, 2018; Habibpour *et al*, 2024; Femerling *et al*, 2022; Galvão & de Lorenzo, 2006). Most bacterial TFs contain small molecule binding domains that enable them to directly sense these intracellular input signals (Ledezma-Tejeida *et al*, 2021; Ziegler & Freddolino, 2021), which may include intermediates of metabolic pathways or imported metabolites. Upon sensing their effector molecules, TFs undergo conformational changes that influence their DNA-binding capacity - either enhancing or inhibiting it – thereby affecting the expression of their target genes (Browning *et al*, 2019).

Since the foundational discoveries regarding bacterial TF signal-sensing mechanisms over 60 years ago (Jacob & Monod, 1961), extensive research has characterized the regulatory interactions between metabolites and TFs, which serve as input to the gene regulation network, particularly in the model organism *E. coli* (Santos-Zavaleta *et al*, 2019; Femerling *et al*, 2022; Browning *et al*, 2019). As of today, input signals have been identified for about 40% of *E. coli*’s TFs that are predicted to directly sense metabolites (Ledezma-Tejeida *et al*, 2021; Femerling *et al*, 2022). However, this proportion decreases dramatically to below 1-2% in non-model organisms (Dudek & Jahn, 2022). The relatively slow pace of the discovery process is largely due to the time-consuming and low-throughput nature of experiments to identify such interactions. Typically, a TF is first characterized regarding the genes and cellular functions it regulates. The search for input signals generally occurs only after extensive studies that map out the TF targets and characterize their functions, usually by hypothesizing potential signal metabolites from the identified cellular function. These hypotheses are then tested individually using *in vitro* methods, such as DNA-binding assays, to elucidate the regulatory effects of these interactions (Ricca *et al*, 1989; Urano *et al*, 2015; Martí-Arbona *et al*, 2012; Nemoto *et al*, 2012; Lim *et al*, 1987; Quail *et al*, 1994; Hart & Blumenthal, 2011; Chavarría *et al*, 2014; Arce-Rodríguez *et al*, 2012).

Recent advances in high-throughput methods now enable the measurement of gene expression and metabolite abundances at large scales, significantly accelerating the identification of TF-metabolite interactions (Holbrook-Smith *et al*, 2024). One effective approach involves correlating *in vivo* TF activities with metabolite abundances. The underlying assumption is that changes in the abundance of an input metabolite for a specific TF should correspond with that TF’s regulatory activity. Although recent pioneering studies have demonstrated this correlation (Kochanowski *et al*, 2017; Lempp *et al*, 2019; Yogendra & Kushalappa, 2016), such applications have thus far been limited to relatively small numbers of TFs or specific growth conditions. Here, we combined the large, publicly available PRECISE2.0 transcriptomics dataset (Rychel *et al*, 2021) with high-throughput metabolomics to systematically predict TF-metabolite interactions in *E. coli*. By quantifying the intracellular metabolome across 40 growth conditions that matched those of the transcriptomics dataset, we obtained paired profiles of gene expression and metabolite abundances across a diverse range of growth conditions. After inferring TF activities from gene expression by leveraging the known *E. coli* regulatory network, we identified correlating pairs of TFs and metabolites, successfully recovering known interactions and revealing novel ones.

## Results

### Inferring transcription factors activity from gene expression

To determine TFs activities, we leveraged published transcriptomics data from the PRECISE2.0 *E. coli* dataset that covers approximately 400 growth conditions (Figure 1a,b) (Rychel *et al*, 2021), as well as known TF target genes from the RegulonDB regulatory network (Figure 1c, Table S1) (Santos-Zavaleta *et al*, 2019). For each TF, activity was inferred based on the expression patterns of its reported target genes, collectively known as its regulon. We tested six different computational methods to assess their ability to correctly predict expected decreases of TF activity in mutant strains from the PRECISE2.0 dataset where a given TF had been knocked out. Among these, the VIPER method (Alvarez *et al*, 2016), which relies on a probabilistic framework to compute enrichment of specific TF regulons in differentially expressed genes, yielded the best results, correctly assigning 34 out of 40 pairs (Figure S1, Methods) and was selected for further analysis. Subsequently, we evaluated whether the inferred TF activities corresponded with the presence of TF input signals. The PRECISE2.0 dataset includes 47 conditions related to 23 TFs, wherein at least one metabolite known to interact with a TF was added to the growth medium. In 83% of these cases, we observed the expected direction of TF activity change when compared to a paired control condition (Figure 1d). These expected changes were evident at both the TF activity and target gene expression levels, as illustrated for four TFs exhibiting diverse combinations of activating/inactivating TF-metabolite interactions and activating/repressing TF-gene interactions (Figure 1e,f). Based on these analyses, we conclude that the inferred TF activities are biologically meaningful and can be effectively used to predict TF input signals.

**Figure 1.**
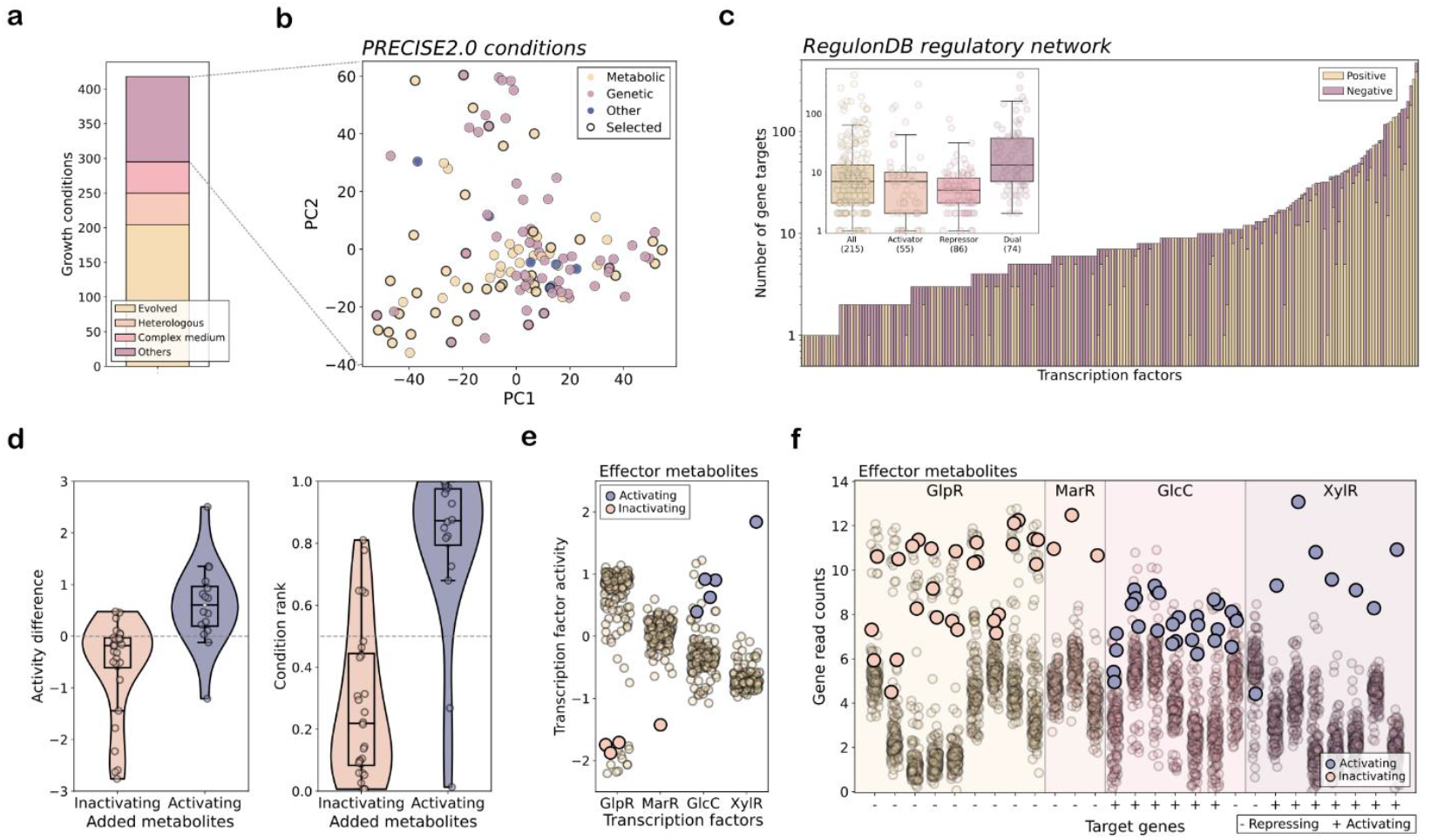
TF activities inferred from gene expression capture the expected effects of TF input signals. (**a**) Distribution of various types of experimental conditions within the iModulon *E. coli* PRECISE2.0 dataset. ‘Evolved’: condition with strain from adaptive laboratory evolution experiment. ‘Heterologous’: condition with strain carrying heterologous DNA. (**b**) Principal component analysis of the candidate experimental conditions in the PRECISE2.0 transcriptomics data, with conditions selected for this study highlighted as black circles. Perturbation types are colored as indicated. (**c**) Enumeration and categorization of regulatory interactions associated with all transcription factors target genes as documented in RegulonDB. (**d**) Impact of adding TF ligand to the medium on the activity of the ligand-regulated TF, presented for all conditions in the PRECISE2.0 dataset where a known TF effector metabolite was added to the growth medium. The effects are illustrated as the activity difference between the TF knockout mutant and the corresponding wild-type control conditions (left), or as the rank of TF mutant condition compared to all other conditions (right). (**e-f**) Influence of ligand addition on TF activity (**e**) and expression of target genes (**f**) for GlpR, MarR, GlcC, and XylR across all conditions. Conditions involving effectors known to activate or inhibit TF activities are indicated in blue or pink, respectively. The direction of known TF regulations for each target gene is shown as ‘-’ for repressive regulation and ‘+’ for activating regulation.

### Selecting a set of reproducible growth conditions

To systematically identify novel TF input signals, we aimed to generate metabolomics data under conditions where TF activities had been inferred, with the goal of finding metabolite abundances that correlate with these TF activities. Approximately 75% of the growth conditions in the PRECISE2.0 dataset pertain to either strains specifically evolved towards phenotypes of interest, strains expressing heterologous DNA, or undefined growth media (Figure 1a). From the remaining 100 conditions, we successfully reproduced the growth condition for 10 strains in 40 cases (Table S2), which will henceforth be referred to as the selected experimental conditions. These selected conditions encompass a diverse array of nutritional and genetic perturbations, including 8 TF knockout mutants (Figure 1b).

For the entire PRECISE2.0 dataset, we obtained a wide range of activities for 173 TFs, highlighting the richness of the transcriptomics dataset (Figure 2a, Table S3). To assess the extent of variability lost by selecting this subset of 40 experimental conditions, we determined the maximum ranges of TF activities; specifically, we quantified the difference between the highest and lowest activity values for each TF across conditions. A higher range value is anticipated to enhance the likelihood of detecting meaningful patterns in the data, as illustrated by the significant increase in correctly assigned TF activities when applying a threshold for higher activity ranges (Figure S1). Overall, the 40 selected experimental conditions, representing about 10% of the PRECISE2.0 dataset, retained an average of 72% of the maximum range values across the entire dataset for the 173 measured TFs (Figure 2b, Table S4).

**Figure 2.**
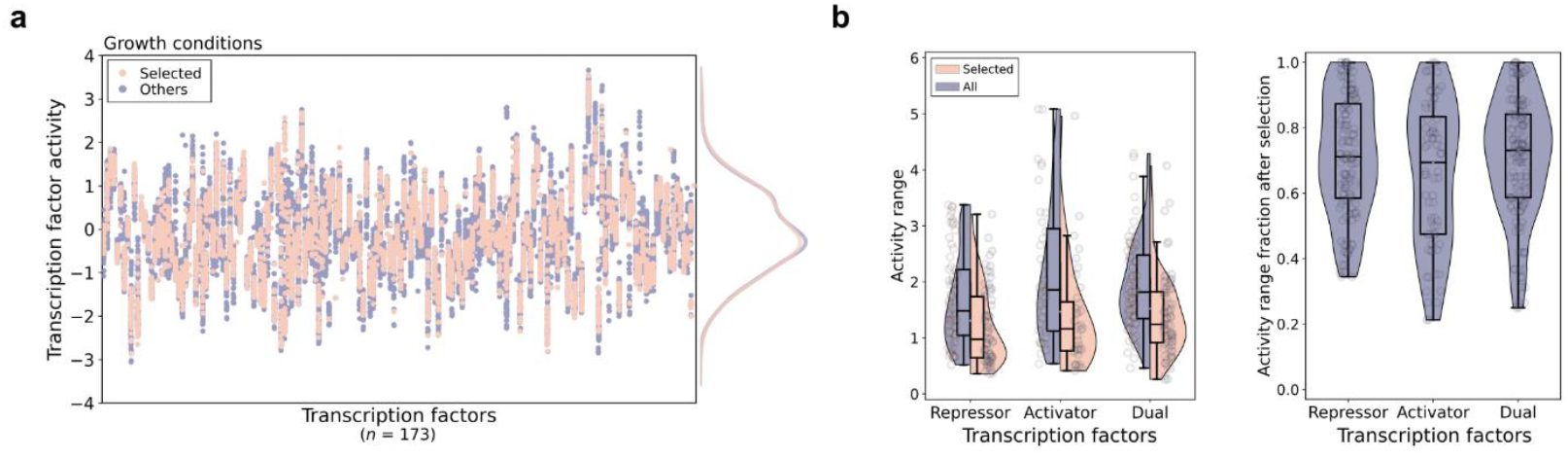
Selected growth conditions preserve data variability. (**a**) Activities of TFs across all experimental conditions in the PRECISE2.0 dataset, encompassing all TFs in RegulonDB with more than 3 known target genes. The selected conditions for which metabolomics measurements were conducted are highlighted in orange and the others are indicated in blue. The density of all TF activities is shown for selected (orange) or all (blue) conditions on the right. (**b**) Left: Activity ranges (difference between highest and lowest activity values) of TFs across either all conditions or the selected ones. Right: Fraction of activity ranges preserved when considering only the selected conditions. TFs are grouped by their gene regulation type: repressor, activator, or dual regulator.

### Systematic prediction of TF-metabolite interactions

To identify novel TF input signals, we cultured *E. coli* wild-types MG1655 and BW25113, as well as eight mutant strains under different growth conditions, corresponding to the 40 selected PRECISE2.0 experimental conditions (Table S2). Intracellular metabolites were extracted during the mid-exponential growth phase, and their abundances were determined using untargeted direct flow-injection mass spectrometry metabolomics (Figure 3a) (Fuhrer *et al*, 2011). As expected, metabolites that were added in specific growth conditions - including known TF signal molecules - exhibited the highest abundance in their corresponding condition (Figure 3b). By utilizing the KEGG database (Kanehisa *et al*, 2017), we annotated 279 metabolites based on their exact masses, for which we obtained relative abundances across a wide range of intensities throughout the 40 selected conditions (Figure 3c, Tables S5, S6). To assess the diversity of metabolic states across these 40 selected conditions, we calculated the maximum abundance fold change for each metabolite. For the 279 annotated metabolites, we found a median abundance fold change of 3.34, indicating significant variation for most compounds and metabolic pathways across the selected conditions. This allowed us to generate a diverse metabolomics dataset aligned with the corresponding published transcriptomics dataset. By correlating metabolite abundances with TFs activities, we were able to generate hypotheses regarding potential metabolic input signals of TFs (Figure 4a).

**Figure 3.**
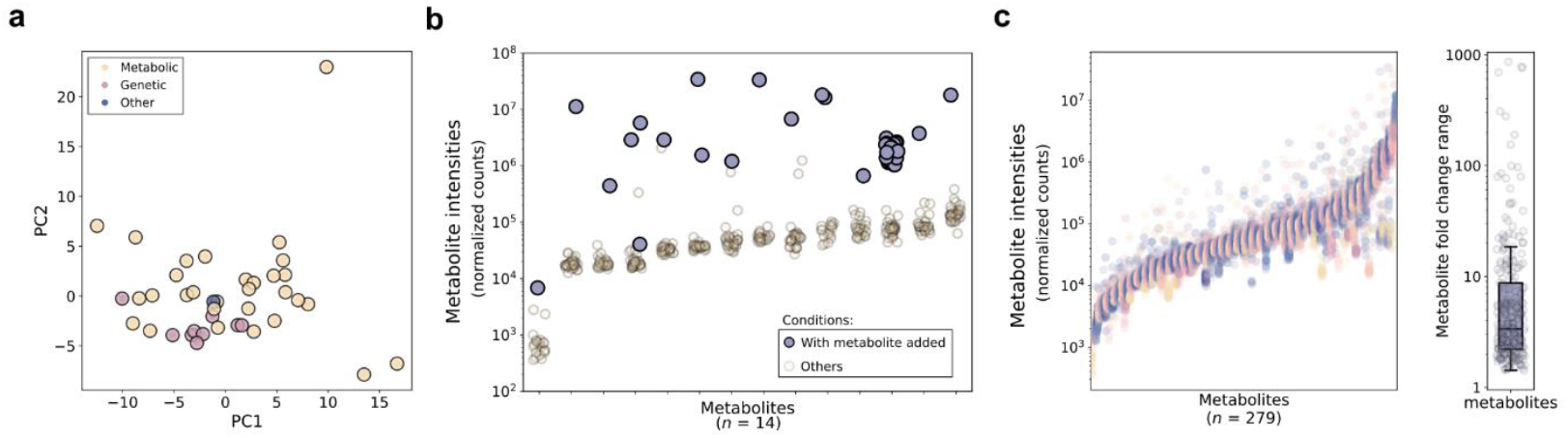
High-throughput metabolomics probes wide ranges of metabolite abundances across diverse growth conditions. (**a**) Principal component analysis of metabolomics data derived from the 40 selected experimental conditions. Perturbation types are colored as indicated. (**b**) Metabolite intensities for 14 metabolites (x axis) that have been added to the growth medium in at least one experimental condition, displayed across all experimental conditions. For each metabolite, the measured intensities are represented by individual dots, with the condition(s) in which it was added marked as blue dots. (**c**) Left: All annotated metabolite intensities across all measured conditions. Different colors are used for different metabolites. Right: Distribution of maximum abundance ranges for all annotated metabolites across all growth conditions.

**Figure 4.**
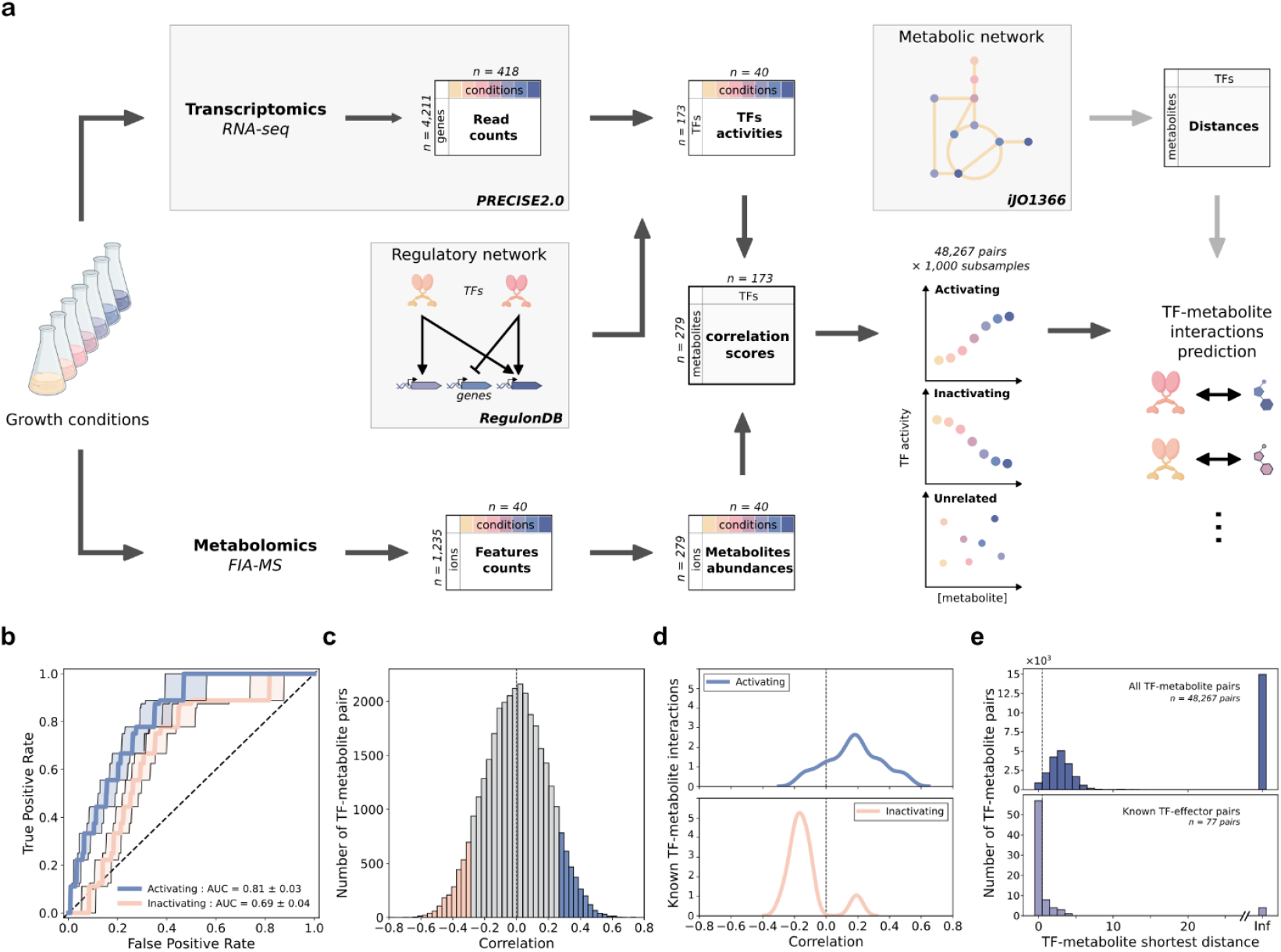
Correlation analysis of TF activities and metabolite abundances recover previously known TF-metabolite interactions. (**a**) Schematic representation of the approach used in this study. The RNA-seq data and the regulatory network were sourced from the iModulon and RegulonDB databases, respectively. We reproduced 40 growth conditions from the iModulon PRECISE2.0 dataset to conduct metabolomics and obtain paired transcriptomics-metabolomics datasets. TF activities were inferred from the transcriptomics data and analyzed in conjunction with metabolite abundances through correlation analysis. (**b**) Median receiver operating characteristic curves illustrating the recovery of known activating (blue) or inactivating (pink) TF-signal interactions. (**c**) Distribution of correlation scores across all TF-metabolite pairs, with bins above or below the false positive rate of 0.1 correlation thresholds in either direction highlighted in color. (**d**) Distribution of correlation scores for known activating (top) or inactivating (bottom) TF-signal interactions. (**e**) Distribution of TF-metabolite distances for known interactions (top) compared to all pairs (bottom).

To ensure the accuracy of our approach and mitigate potential biases stemming from single outliers or poorly matching growth conditions, we used bootstrapping techniques on the selected conditions for correlation. Specifically, we randomly sub-sampled 30 out of the 40 selected conditions across 1,000 samples and assessed the performance of our correlation approach for each sample. Notably, most previously known TF-metabolite interactions were amongst the top correlating pairs (Figure 4b), illustrating the efficacy of our approach in identifying TFs input signals. Interactions that increase TF activity were particularly well-recovered with an area under the receiver operating characteristic (ROC) curve of 0.81. Conversely, the recovery of interactions that decrease TF activity showed lower performance in negatively correlating pairs at low false positive rates (Figure 4b); thus, we focused on activating interactions moving forward. In total, we obtained average correlation values for 48,267 TF-metabolite pairs across the 173 TFs (Figure 4c,d), with highly correlating pairs suggesting numerous potential new interactions.

To generate a list of higher-confidence candidate TF input signals, we implemented three filtering methods to assess the likelihood of TF-metabolite pairs being involved in regulatory interactions (Figure S2). First, we applied a false positive rate threshold of 0.1 based on the median ROC curves. Second, we established a stability metric that indicates, for each pair, the proportion of bootstrapping samples in which the correlation score exceeds the false positive rate threshold. The stability threshold was then defined as the value that maximizes the retention of known TF-metabolite interactions. Third, we defined a distance metric representing the number of consecutive reactions in the metabolic network needed to connect the metabolite involved to the nearest enzyme directly regulated by the TF. Setting a threshold for lower distance values increases the likelihood of identifying functional pairs (Figure 4e) (Lempp *et al*, 2019), because TF input signals typically originate from within or near the regulated pathways (Santos-Zavaleta *et al*, 2019). However, this filtering restricted our search to TFs with at least one regulated enzyme, which included 132 out of the 173 TFs with inferred activities. For positively correlating pairs, applying the stability and distance filters increased the proportion of recovered known interactions among predicted pairs about 100-fold, resulting in a final list of 103 TF-metabolite pairs, only 5 of which were previously known (Figure 5a, Table S7). Overall, this approach identified candidate input signals for 42 TFs.

**Figure 5.**
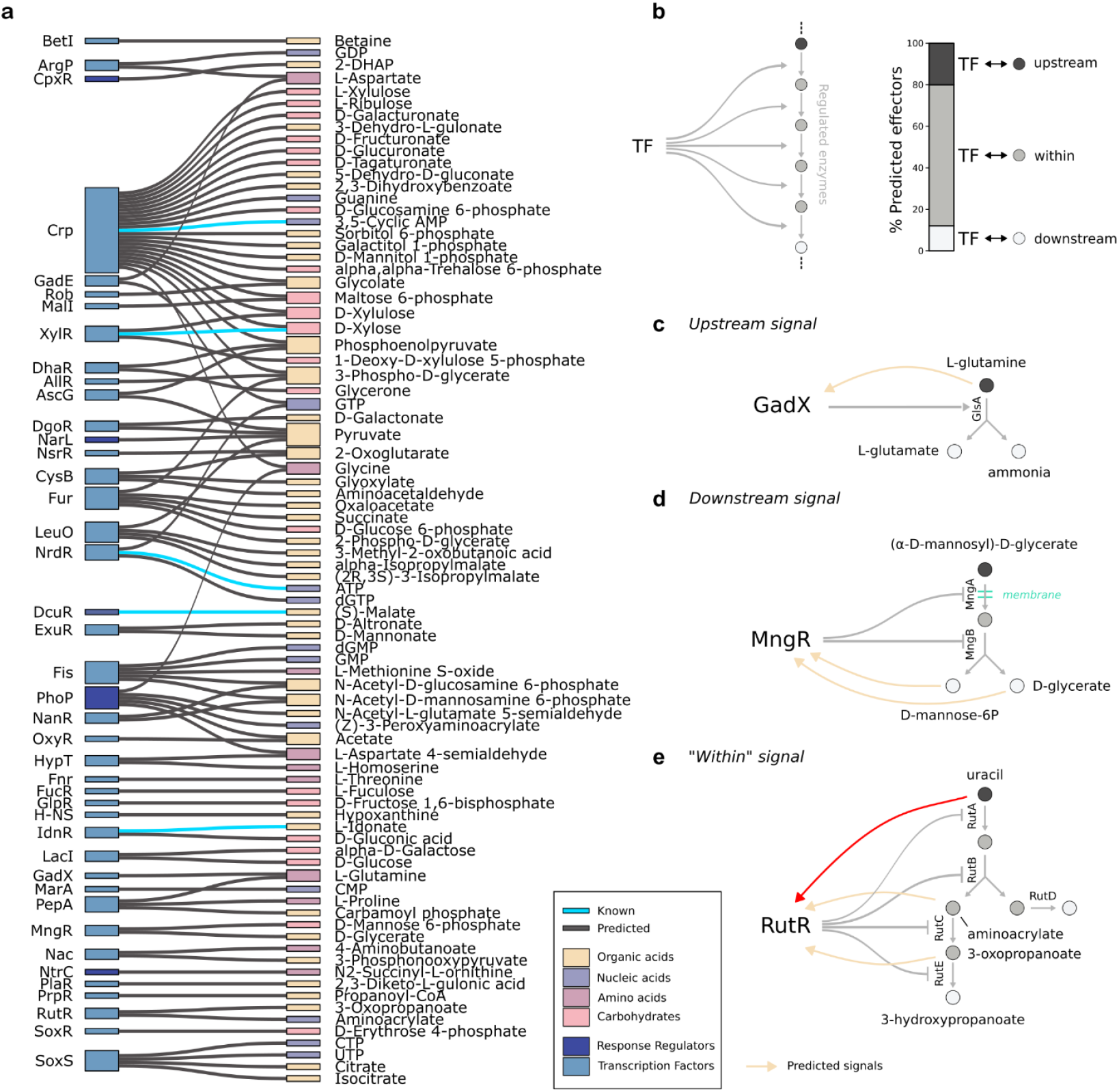
Predicted input signals encompass a wide range of chemical classes and regulatory logics. (**a**) Sankey diagram illustrating the predicted pairs of TFs and signals. TFs and metabolites are color-coded according to different categories, as detailed in the graphical legend. Previously known activating TF-metabolite interactions are indicated with light blue links. (**b**) Proportions of predicted signals based on their location within the metabolic sub-network that is directly regulated by the respective TF. (**c-e**) Example predictions for upstream (**c**), downstream (**d**), or within (**e**) predicted metabolic input signals. Predicted TF-signal interactions are represented with yellow arrows. Metabolites are colored as shown in panel b, while regulated enzymatic reactions are depicted in grey. For RutR (**e**), the previously known inactivating input signal is shown as a red arrow.

Among the 42 TFs with predictions, 18 had previously known small molecule input signal, including eight that enhanced TF activity and were detected by our metabolomics analysis (Table 1). Our approach correctly predicted the known signal molecules in five out of these eight cases. For the remaining three cases, the predicted metabolite was found directly downstream of the known signal, being the product of an enzymatic reaction in which the known signal serves as the substrate. A potential explanation for this could be rapid conversion of the actual signal molecule during sample processing. Altogether, our approach successfully recovered five of the eight known signal molecules for both direct and indirect TF activations, while in the other cases it predicted a metabolite one reaction step removed from the signal.

Among the predicted pairs, Crp was the TF with the highest number of candidate effectors, totaling 23 candidates, while the average across of TFs was below two (Table S7). With over 600 known targets, Crp possesses the largest array of target genes among all *E. coli* TFs, including hundreds of enzymes. Consequently, many metabolites are in close proximity to Crp targets, which diminishes the effectiveness of our distance filtering process. Additionally, the number of predictions was inflated due to the inability of our metabolomic approach to distinguish several metabolites with the same mass, particularly among the predicted sugar inputs to Crp. Despite the relatively high number of candidates generated for Crp, its known effector, cyclic AMP, was among the predicted candidates and had the highest correlation and stability scores.

Beyond Crp, we predicted 80 candidate effector metabolites for 41 TFs. These predicted pairs represent all three established regulatory patterns (Santos-Zavaleta *et al*, 2019) (Figure 5b). Upstream: the metabolite is the first substrate in a metabolic reaction, or a series of reactions regulated by the TF. Within: the metabolite is an intermediate in a chain of reactions regulated by the TF. Downstream: the metabolite is an end product of a reaction, or a series of reactions regulated by the TF. Novel pairs were predicted for each regulatory pattern with the majority of pairs being from the within pattern (Figure 5b), which is consistent with known interactions (Santos-Zavaleta *et al*, 2019). 12 TFs had upstream metabolites identified as input signals, indicating feedforward sensing of the substrates of the regulated reactions. For example, L-glutamine was predicted to be the activating signal of GadX. GadX controls the acid resistance response notably with the activation of *glsA* (Tucker *et al*, 2003; Tramonti *et al*, 2008), which is involved in acid resistance through the conversion of L-glutamine to L-glutamate and ammonia (Lu *et al*, 2013). As a predicted activator of GadX, L-glutamine thus promotes the positive regulation of that pathway (Figure 5c). Nine cases involved metabolites that were downstream of target enzymes. An example is the repressor MngR, which inhibits two proteins engaged in uptake and cleavage of 2-O-alpha-mannosyl-D-glycerate (Figure 5d) (Sampaio *et al*, 2004). In this instance, both cleavage products, mannose-6-P and glycerate, were predicted to activate the inhibiting TF, representing a negative feedback loop. Lastly, 27 TFs were associated with predicted signals that correspond to metabolites within their regulated pathways, such as RutR (Figure 5e), which regulates pyrimidines metabolism (Shimada *et al*, 2008) and has two previously known inactivating input signals: uracil and thymine (Shimada *et al*, 2007). Aminoacrylate and 3-oxopropanoate were both predicted as activating signals of RutR from within the RutR-repressed uracil degradation pathway (Figure 5e). While uracil induces upregulation of the pathway from upstream through the inactivation of RutR, the two predicted signals located after the branching out from carbamate represent negative feedback loops to inhibit the pathway from within that end branch. This negative regulation of the pathway could be needed in case of accumulation of 3-oxopropanoate, which is toxic (Parales & Ingraham, 2010). Overall, we identified 76 novel candidate input signals spanning all previously known regulatory logics.

### Validation of predicted input signal for LeuO

For some TFs, we predicted multiple plausible input signals with varying regulatory logics. A notable example is LeuO, the activator of the leucine biosynthesis operon that is also a global antagonist against the universal silencer of stress-response genes H-NS (Shimada *et al*, 2011; Sánchez-Popoca *et al*, 2022), for which no input signal was previously known (Fragel *et al*, 2019; Santos-Zavaleta *et al*, 2019). We predicted four LeuO input signals derived from either upstream or within the leucine biosynthesis pathway (Figure 6a). The prediction of these multiple input signals may stem from consecutive metabolites displaying correlated abundances, leading to similarly high scores in our analysis. In such scenarios, experimental validation is crucial to pinpoint the exact signal. We chose LeuO for this validation to determine whether one of its four predicted signals influences its DNA-binding function. The LeuO protein was purified and its binding to DNA assessed at the promoter region of one of its target operons, *leuLABCD*, using an electro-mobility shift assay. To maximize the dynamic range of our assay for measuring the effect of an added molecule on LeuO’s DNA-binding activity (either up or down), we first established a LeuO concentration where approximately half of the DNA fragments were bound by the TF, as determined by a concentration gradient (Figure S3). Using these conditions, we tested the effect of each of the four predicted input signals on LeuO activity and found that only 2-isopropylmalate significantly enhanced LeuO binding to its target DNA (Figure 6b, Figure S4). To further confirm this novel interaction, we employed a thermal shift assay to assess the influence of 2-isopropylmalate on the stability of the LeuO-DNA complex. Typically, an input molecule modifies the conformation of its bound TF, which in turn impacts the stability of the TF-DNA complex. In this assay, 2-isopropylmalate significantly increased the thermal stability of LeuO by 1.6°C (Figure 6c), thereby further validating 2-isopropylmalate as the input signal to LeuO.

**Figure 6.**
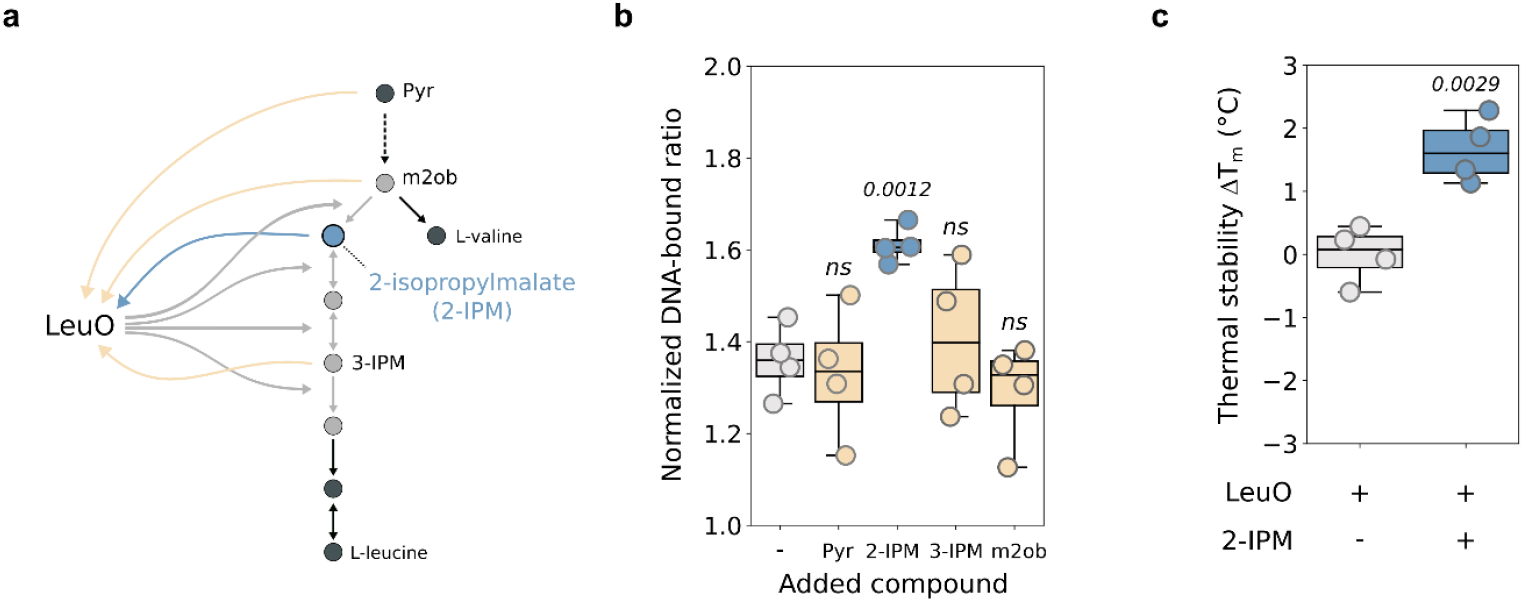
LeuO responds to 2-isopropylmalate. (**a**) Predicted activating input signals for LeuO. Predicted TF-signal interactions are shown by yellow or blue arrows. The blue arrow represents the signal metabolite (2-isopropylmalate) found to interact with LeuO in **b,c**. Metabolites are colored as in Figure 5b. Regulated enzymatic reactions are shown in grey. The dotted reaction arrow represents multiple consecutive enzymatic reactions not shown on the scheme. Pyr: Pyruvate, m2ob: 3-methyl-2-oxobutanoate, 3-IPM: 3-isopropylmalate. (**b**) Change in LeuO DNA-binding activity on the *leuLABCD* promoter region upon addition of predicted signal metabolites measured by electromobility shift assay. Predicted signal metabolites are named with letters (a to d) following the annotation in **a**. DNA binding is measured as the ratio of bound to unbound DNA fragments for each gel lane normalized to a control lane without LeuO protein on each gel from four independent gel shift experiments (individual dots). (**c**) Change in thermal stability of the LeuO-DNA complex upon addition of 2-isopropylmalate (2-IPM) measured by thermal shift assay in four independent experiments (individual dots). Results are shown for each replicate as the difference between its melting temperature and the average melting temperature of the control samples without 2-IPM (ΔT_m_). Exact *p*-values below 0.05 are written above the corresponding box when compared to the control samples. ns: non-significant.

## Discussion

Even in *E. coli*, arguably the best-studied bacterium, we still do not know the signals that most TFs sense to initiate transcriptional responses (Femerling *et al*, 2022; Ledezma-Tejeida *et al*, 2021). This knowledge gap can primarily be attributed to the labor-intensive and low-throughput methodologies that have been employed to elucidate these interactions. Using paired transcriptomics and metabolomics, we correlated the regulatory activities of TFs with the abundance of metabolites to predict signal molecules across 173 TFs. Our approach recovered most known activating TF-metabolite interactions, among the detected metabolites, but also unveiled numerous novel interactions. Upon refining our predictions, we identified a high-confidence ensemble of predicted TF-metabolite interactions encompassing 80 metabolites and 41 TFs. These predicted signal molecules span a wide range of chemical classes and regulatory patterns, including both metabolic feedforward and feedback regulations. The outcome of this study proposes a reproducible experimental pipeline to identify TFs input signal with unprecedented throughput. This advancement identified numerous novel regulatory interactions and brings us closer to a comprehensive transcriptional regulatory network in *E. coli*.

For experimental validation, we focused on LeuO, a TF that regulates diverse cellular processes in gammaproteobacteria (Hernández-Lucas & Calva, 2012), including its most conserved target: the adjacent leucine biosynthesis operon *leuABCD*. As a LysR-like TF, LeuO features a predicted metabolite-binding domain. Notably, mutations that mimic a bound state enhance LeuO’s regulatory activity; however, the input signal remained unidentified (Fragel *et al*, 2019). We experimentally validated the predicted 2-isopropylmalate as the signal metabolite that increases LeuO activity upon binding. As an intermediate in leucine biosynthesis, 2-isopropylmalate is the first pathway-specific metabolite to emerge after the divergence from valine biosynthesis (Calvo *et al*, 1962). Our findings suggest a positive feedforward loop whereby LeuO activation promotes leucine biosynthesis in response to the accumulation of this initial pathway-specific metabolite. While our results clarify regulation of the leucine pathway, the regulatory logic by which LeuO governs processes beyond leucine synthesis remain uncertain, especially considering its more than 100 gene targets (Dillon *et al*, 2012; Shimada *et al*, 2011; Hernández-Lucas & Calva, 2012). This scenario bears resemblance to other broadly acting TFs, such as Lrp, PdhR or ArgP, which also respond to a single biosynthesis pathway-specific metabolite (Kroner *et al*, 2019; Anzai *et al*, 2020; Nguyen Le Minh *et al*, 2018). In these cases, the pathway related to the input signals are typically well conserved among the target genes, while additional targets display variability across species or strains, indicating their later incorporation into the TF regulon (Trouillon *et al*, 2020; Baumgart *et al*, 2021).

While the approach presented here accelerates the identification of TFs input signals, it has several limitations. Firstly, the activities of TFs are inferred from previously identified gene targets for each TF, which means our approach heavily relies on existing knowledge of the transcriptional network (Badia-i-Mompel *et al*, 2022). This knowledge base is currently quite incomplete for most organisms (Santos-Zavaleta *et al*, 2019), even in *E. coli* covering only about 30% of the regulatory network (Trouillon *et al*, 2023). Consequently, predictions could be made for only 173 out of approximately 300 predicted TFs in *E. coli*. Expanding the knowledge base not only increases the number of TFs with predictions but would also enhance the quality of these predictions by refining and enlarging the set of TF-regulated targets. Even well-studied TFs often lack comprehensive target annotations or may have reproducibility issues with their annotated targets. Secondly, the selection and number of experimental conditions used for correlation analysis strongly influences the potential for identification of new interactions (Lamoureux *et al*, 2021). In a previous study, the analysis was limited to transitions between two growth conditions, which restricted the interpretation of results, as many metabolites exhibited similar abundance dynamics (Lempp *et al*, 2019). We mitigated this issue by measuring metabolite abundances across 40 diverse experimental conditions, thereby reducing the likelihood of obtaining highly similar profiles. Nevertheless, the specific choice of conditions remains influential, as many TFs are only expressed or active under particular circumstances. Lastly, the number and types of metabolites measured are contingent upon the chosen metabolomics method (Alseekh *et al*, 2021). By relying on MS1-based annotation, several metabolites could not be distinguished in our analysis, potentially resulting in an artificially inflated number of predicted metabolites in some cases, such as for Crp. To enhance coverage of detected metabolites, the integrated use of multiple metabolomics methods may prove beneficial.

The application of high-throughput inference methods often results in elevated false positive rates compared to targeted *in vitro* validation experiments (Lempp *et al*, 2019; Piazza *et al*, 2018). In scenarios where multiple candidate signals are predicted - such as with LeuO in this study - experimental validation is essential for accurately identifying the specific signal. Techniques like thermal shift assays and microscale thermophoresis are capable of detecting TF-metabolite interactions (Huynh & Partch, 2015; Jerabek-Willemsen *et al*, 2014); however, it is important to note that these interactions do not always correlate with functional effects, necessitating further characterization. Currently, the gold standard for functional validation are electro-mobility shift assays (Hellman & Fried, 2007), which, despite their reliability, are labor-intensive and impractical for large-scale applications. Consequently, we concentrated on one of our prediction cases, LeuO, and performed a comprehensive validation using two complementary methods. The functional validation of these interactions constitutes a significant bottleneck in achieving a fully mapped regulatory network. Given that inference methods like the one presented here can generate hypotheses for hundreds of TFs, there is a pressing need for a functional validation method that can match this throughput.

Collectively, these results demonstrate the potential of high-throughput approaches for the rapid discovery of TF-metabolite interactions, revealing numerous such interactions in *E. coli*. By facilitating the generation of hypotheses on TF input signals at the scale of hundreds of TFs, we anticipate that approaches like the one presented here will significantly accelerate advancements the field of TF-metabolite interaction discovery.

## Methods

### Bacterial strains and growth

Bacterial strains, media composition and growth conditions were chosen to match experimental procedures used to generate RNA-seq data in iModulonDB (Lamoureux *et al*, 2021) (Table S2). All samples were cultivated in biological triplicates in 14 ml tubes (Greiner, #187262) at 37°C and 250 rpm agitation, in 3 ml of M9 minimal medium (42.2 mM Na_2_HPO_4,_ 22 mM KH_2_PO_4_, 18.7 mM NH_4_Cl, 8.5 mM NaCl, 1 mM MgSO_4_, 0.1 mM CaCl_2_, 20 µM FeCl_3_, 2.4 µM ZnCl_2_, 2.1 µM CoCl_2_, 2.1 µM Na_2_MoO_4_, 1.3 µM CuCl_2_, 2 µM H_3_BO_3_) supplemented with different carbon sources and supplements prior to growth experiments. For protein expression, strains from the ASKA library (Kitagawa *et al*, 2005) were grown in lysogeny broth medium containing 30 µg/ml chloramphenicol.

### Metabolites extraction

After overnight culture in M9 medium, cells were diluted to OD_600_ = 0.05 and cultivated until mid-to late-exponential phase to match the harvest OD_600_ reported for RNA extraction. Metabolites were then extracted by fast filtration. Briefly, a volume of culture corresponding to a 1 ml equivalent of OD_600_ = 1 was vacuum-filtered on a 0.45 μm nitrocellulose filter (Durapore, #HVLP04700). Immediately after filtration, metabolic processes were quenched by putting the filter membrane into 1 ml of extraction solution (40% (v/v) acetonitrile, 40% methanol, 20% ddH2O) precooled at -20°C in 6-well plates. Plates were sealed with parafilm and metabolites were extracted overnight at −20°C. After centrifugation of the extraction solution for 8 min at −2°C and 21,000 g, 650 μl of supernatant were collected and stored at −80°C until measurement.

### Metabolomics

Extracts were measured in technical duplicates by double injection on an Agilent 6550 quadrupole time-of-flight mass spectrometer coupled to a Gerstel MPS2 autosampler using FIA-TOF-MS, as previously described (Fuhrer *et al*, 2011). 5 μl of sample were injected into a constant 150 µl/min flow of running buffer (60:40 (v/v) isopropanol:water, 5mM ammonium carbonate at pH 9, containing 3-amino-1-propanesulfonic acid and hexakis(1H, 1H, 3H-tetrafluoropropoxy)phosphazine for online mass axis correction). Mass spectra were recorded in negative ionization and high resolution modes with an acquisition rate of 1.4 spectra/s. Mass spectrometry data was merged, normalized and annotated in Matlab R2021a. After spectral merging, counts were normalized by total ion count. Then, measured ions were annotated by mass-to-charge ratios to a reference list derived from a genome-wide reconstruction of *E. coli* metabolism within 0.001 Da mass tolerance. Biological and technical replicates were filtered for a coefficient of variation smaller than 15 %.

### Estimation of transcription factors activities

RNA-seq data from the PRECISE 2.0 dataset of iModulonDB (Lamoureux *et al*, 2021) and the *E. coli* transcriptional regulatory network from RegulonDB v 10.5 curated from iModulonDB (Santos-Zavaleta *et al*, 2019) were used to infer transcription factors activities using the decoupleR python package v 1.2.0 (Badia-i-Mompel *et al*, 2022) on TFs with at least three known target genes. To account for incompleteness of the regulatory network, the network was subsampled, as previously described (Ortmayr *et al*, 2019), for each TF within ten subnetworks containing 40 randomly chosen additional TFs for which activities were inferred. For each TF, the median of all subsampling simulations was calculated and used as final activity value.

To assess the performance of different inference methods, conditions from iModulonDB (Lamoureux *et al*, 2021) that had single TF gene knockouts and matching wild-type strains were selected to compare inferred changes to expected directions; i.e., decreased TF activity in the corresponding TF mutant strain. Six methods from the decoupleR package were compared (Badia-i-Mompel *et al*, 2022) using a direction metric and median rank percentile metric(Ma & Brent, 2021). As a robust estimate, the median and interquantile range across all subsamples per TF in each condition was calculated. Accuracy was reported for each TF as the proportion of samples in which the activity is lower in the TF mutant compared to wild-type strain (Figure S1). Based on accuracy on the direction metric, all TFs were filtered on baseline activity and minimum absolute activity difference as thresholding on these criteria showed improved performance on correctly assigned TF activities. To that aim, thresholds were chosen as minimizing the *X*^2^ *p-value* when dynamically thresholding across the range of corresponding values for increased correctly assigned TF activities. The VIPER method was chosen to perform all further TF activity inference as it correctly assigned expected TF activity changes for the most conditions (Figure S1).

### Determination of distances within the regulatory and metabolic networks

To obtain distances between TFs and metabolites, TF-gene interactions were obtained from the RegulonDB v10.5 transcriptional regulatory network (Santos-Zavaleta *et al*, 2019) and metabolic reactions from the iJO1366 *E. coli* metabolic model (Orth *et al*, 2011). All highly connected metabolites (involved in >50 reactions) and common cofactors were removed before calculation and only intracellular metabolites and reactions were considered. For each TF-metabolite pair, a possible path between the two networks was searched if at least one of the target genes of the involved TF encodes an enzyme catalyzing a reaction present in the metabolic model. The shortest possible path was determined using the Dijkstra algorithm (Dijkstra, 2022). For a given TF-metabolite pair, if the metabolite is a product or a substrate of an enzyme whose gene is regulated by the involved TF, the assigned distance was zero. The distance was increased by one for each linked reaction needed to reach the closest regulated enzyme. If no path was found, the pair got assigned an infinite distance value. To determine the positions of metabolites within regulated subnetworks (i.e. downstream, upstream or within), the directions of reaction were obtained from the iJO1366 *E. coli* metabolic model (Orth *et al*, 2011).

### Inference of TF-metabolite interactions

To predict TF-metabolite interaction candidates, we calculated pairwise Spearman correlations between TF activities and metabolite abundances. To decrease the effect of potential single condition outliers, we used a bootstrapping approach and randomly subsampled 30 out of the 40 growth conditions over 1,000 samples. For each sampling, Spearman’s rank correlation coefficients were calculated between TF activities and metabolite abundances over the 30 conditions and used to generate a receiver-operator characteristics (ROC) curve using reported TF-metabolite interactions as true positives (Santos-Zavaleta *et al*, 2019) and all other pairs as negative cases. Correlation scores are reported for each TF-metabolite pair as the average correlation coefficient for that pair across all 1,000 subsamples. Median ROC curves across all subsamples were used to set a correlation threshold controlling for a 0.1 false positive rate. For each TF-metabolite pair, a stability score was calculated as the proportion of subsamples above the correlation threshold.

To infer potential TF-metabolite interactions, pairs were filtered based on three criteria: their correlation, stability and distance scores. Based on the recovery of true positives, thresholds were determined for each metric (Figure S2) and pairs scoring above all thresholds were selected in the final lists of potential interactions (Table S7). The correlation threshold was chosen to obtain a 0.1 false positive rate. The stability threshold of 0.128 was chosen as the point maximizing the proportion of retained true positives. The distance threshold of 2 was chosen as one above the value retaining 90% of true positives, as a more permissive choice to increase chances of discovering interactions involving metabolites also outside of the regulated enzymatic reactions.

### Protein purification

Selected TFs were purified from overexpressing strains from the ASKA library (Kitagawa *et al*, 2005). Bacteria were grown in LB medium containing 30 µg/ml chloramphenicol. Growth was started from an overnight pre-culture into 100 ml of medium at an OD_600_ of 0.05 at 37°C under agitation (250 rpm). When the OD_600_ reached 1, cultures were cooled to room temperature (RT) and 0.5 mM IPTG was added before resuming growth at RT under agitation for 16 hours. Bacteria were then centrifuged at 3,000 g for 20 min at 4°C. Pellets were resuspended in 10 ml of Lysis buffer (50 mM Tris-HCl, 500 mM NaCl, 10 mM imidazole, 5% glycerol, pH 8) containing freshly added protease inhibitors (Roche, #11836170001) and sonicated at 50% power for 2 cycles of 5 min using a Bandelin Sonopuls HD 2070 sonicator. Lysates were clarified by centrifugation at 3,000 g for 30 min at 4°C. Supernatants were then incubated for 1 h on a rotating wheel with 500 µl of cobalt resin (Thermo, #89964) previously washed three times with Lysis buffer. Samples were then passed through empty gravity columns (Thermo, #29924) and washed once with Lysis buffer, twice with Lysis buffer containing 20 mM imidazole and twice with Lysis buffer containing 40 mM imidazole. After the final wash, columns were closed and 2 ml of Elution buffer (50 mM Tris-HCl, 500 mM NaCl, 200 mM imidazole, 5% glycerol, pH 8) were added. After incubating for 10 min, eluates were collected and then buffer exchanged in storage buffer (50 mM Tris-HCl, 250 mM NaCl, 50 mM KCl, 10% glycerol, 0.1% Tween20, pH 8) using centrifugal filter units (Millipore, #UFC500324). Protein quality was assessed by SDS-PAGE and concentration was measured using a Qubit fluorometer.

### Electro-mobility shift assays

Electro-mobility shift assays were performed as previously described (Trouillon *et al*, 2021). Briefly, Cy5-labeled DNA probes were generated by PCR from *E. coli* MG1655 genomic DNA using specific primers containing a Cy5 modification (Table S8). Binding reactions were performed in EMSA buffer (10 mM Tris-HCl, 50 mM KCl, 5 mM MgCl_2_, 5% glycerol, 0.1 mg/ml BSA, pH 7.5) in 20 µl final volume containing 0 or 10 mM of tested metabolite, 10 nM DNA probe, 500 ng salmon sperm DNA and 2 µl of purified protein, added last. After a 15 min incubation at room temperature, 10 µl of sample were loaded on native 5% TBE gels (Bio-Rad, #4565015) and ran in 0.5× TBE buffer at 120V at 4°C for 1 h. Gels were visualized using a Fusion FX6 Edge imaging system and band intensities were quantified using GelBox (Gulbulak *et al*, 2024).

### Thermal shift assay

Thermal shift assays were performed using the Protein Thermal Shift Dye Kit (Thermo, #4461146) following manufacturer’s instructions. Briefly, 2 µg of LeuO protein, 225 ng of *leuLABCD* promoter region DNA fragment and 0 or 10 mM of 2-isopropylmalate were incubated in 12.5 µl of EMSA buffer (containing no BSA) for 15 min at room temperature. Then 5 µl of Protein Thermal Shift Buffer and 2.5 µl of Protein Thermal Shift Dye were added and melt curves were recorded on a QuantStudio 3 Real-Time PCR System (Applied Biosystems).

## Acknowledgments

We are grateful to members of the lab of Prof. Bernhard O. Palsson, especially Kevin Rychel, Ying Hefner, and Ye Gao, for their help and advice in using data from iModulonDB and reproducing growth conditions. We thank Prof. Mattia Zampieri for his valuable advice and discussions on transcription factor activity inference and the correlation analysis between transcription factor activities and metabolite concentrations. Funding was provided by the ETH Postdoctoral Fellowship program and an ETH Career Seed award (ETH Zürich) for J.T.

## Author contributions

J.T. and U.S. conceptualized the project. J.T., A.E.H. and Y.T. performed experiments and analyzed the data. J.T. and U.S. wrote the manuscript.

