## Supplementary Figures for "Predicting input signals of transcription factors in *Escherichia coli* through metabolomics and transcriptomics"

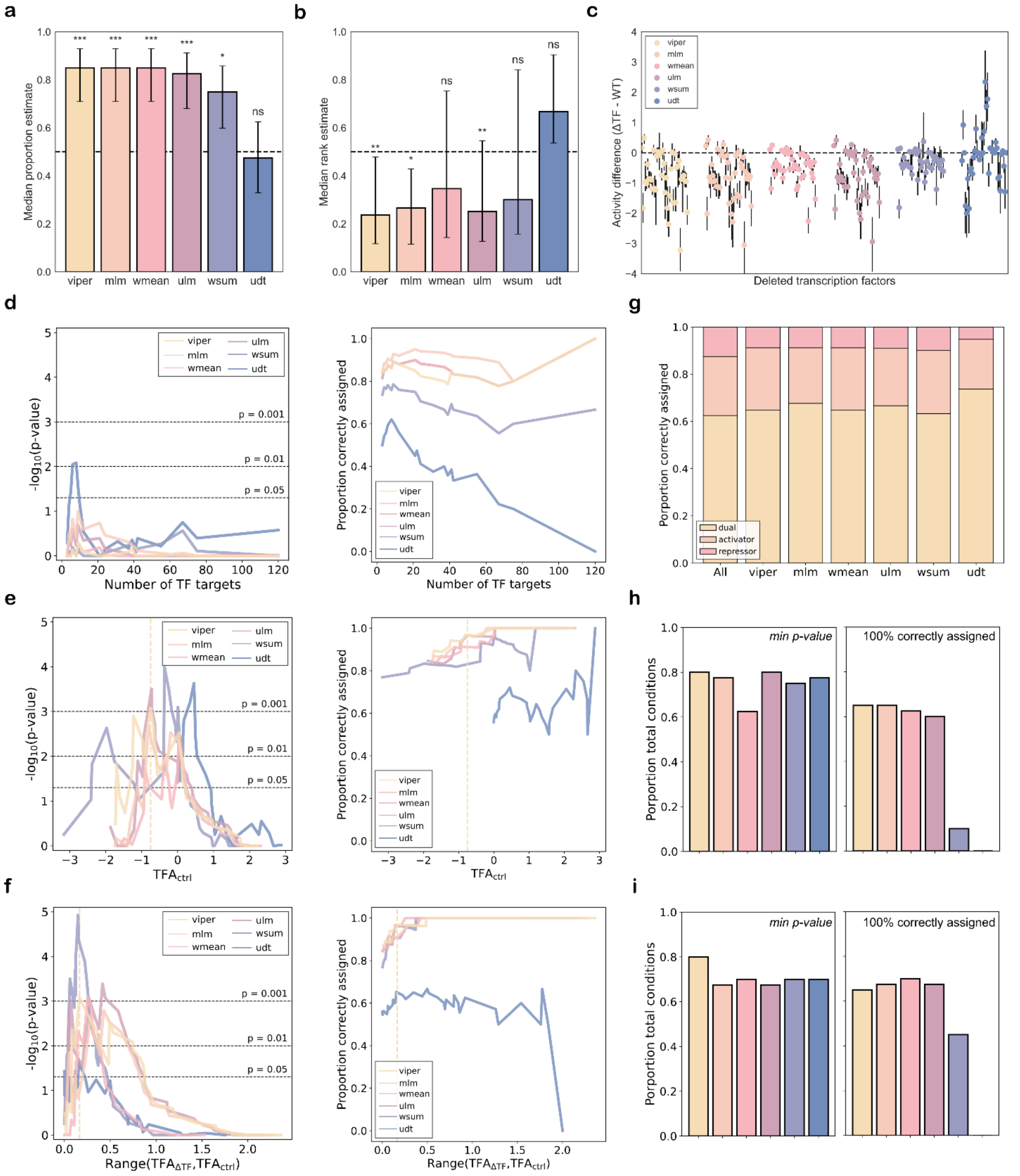


**Figure S1: Validation of transcription factor activity inference methods.** (**a**) Proportion of TF mutant conditions with correct direction inference; i.e., the corresponding TF activity is lower in the TF knockout mutant sample compared to a control. Bars indicate the 95% Wilson score intervals. (**b**) Median rank estimates of all deleted TFs across all TFs in TF mutant conditions. Bars indicate interquantile ranges from TF activities across conditions. (a,b) *ns*: non-significant, *: $p < 0.05$, **: $p < 0.01$, ***: $p < 0.001$ from a two-sided binomial test against random distribution (dashed line) with Bonferroni correction. (**c**) TF activity difference (mutant against corresponding *wild type*) for each TF mutant condition across the six tested methods. (**d,e,f**) Left: Statistical significance from $X^2$ test for increased correctly assigned TF activities resulting from dynamical thresholding across ranges of values of (**d**) number of gene targets of the tested TF, (**e**) TF basal activity in the corresponding control condition or (**f**) TF activity range between the control and TF knockout mutant conditions. TFA: Transcription factor activity. Right: Proportion of correctly assigned TF activities from TF mutant conditions across a range of threshold values based on (**d**) number of gene targets of the tested TF, (**e**) TF basal activity in the corresponding control condition or (**f**) TF activity range between the control and TF mutant conditions. Threshold values minimizing the *p*-value are shown as dotted vertical lines for the viper method when significant. (**g**) Proportion of correctly assigned TF activities from TF mutant conditions split between TF types (activator, repressor or dual regulators). (**h**,**i**) Porportion of experimental conditions kept after thresholding for (**h**) TF basal activity in the corresponding control condition or (**i**) TF activity range between the control and TF mutant conditions using either the threshold values minimizing the *p*-value (left) or the threshold values that allow for 100% of correctly assigned TF activities when possible (right). (a-i) The six tested methods are multivariate linear model (mlm), weighted mean (wmean), virtual inference of protein-activity by enriched regulon analysis (viper), univariate linear model (ulm), weighted sum (wsum) and univariate decision tree (udt). All methods are colored following the same color scheme across all panels.

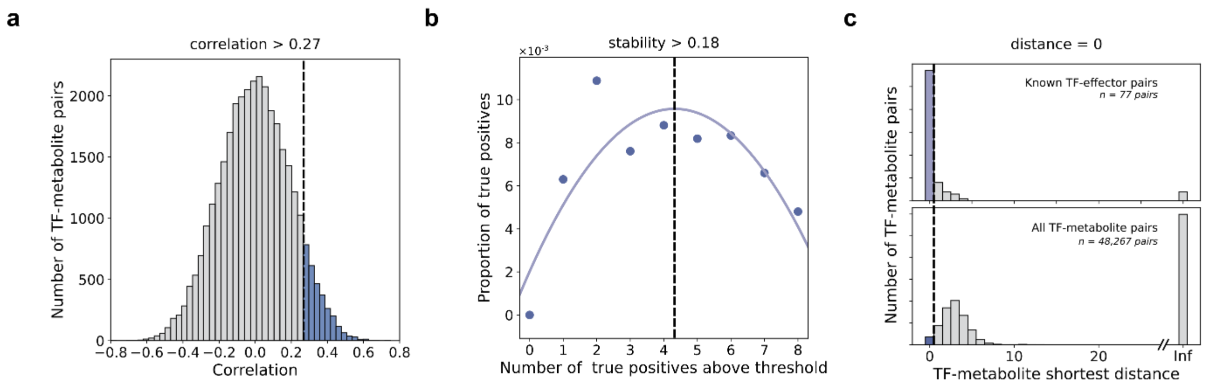


**Figure S2: Filtering approach to identify candidate TF input signals.** (**a**) Distribution of correlation scores across all TF-metabolite pairs. Bins above the False Positive Rate of 0.1 correlation threshold (dotted line) are colored in blue. (**b**) Proportion of known interactions recovered across a range of stability thresholds expressed as number of known interactions above threshold. The stability threshold was determined as the summit of a parabolic fit (dotted line). (**c**) Distribution of TF-metabolite distances for known interactions (top) or all pairs (bottom), with distance threshold of 0 shown as dotted line.

a

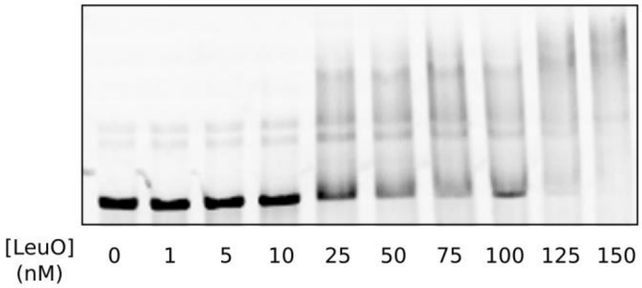


b

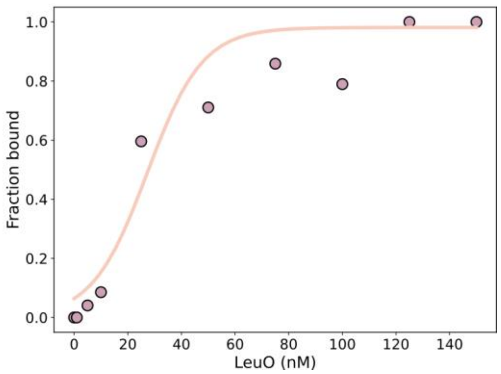


**Figure S3: LeuO binding to the *leuLABCD* promoter region.** (**a**) Gel shift experiment with increasing concentrations of LeuO protein and 0.5 nM of DNA fragment containing the *leuLABCD* promoter region. (**b**) Quantification of fractions of bound DNA from **a**. The curved line represents a sigmoid fit of the data.

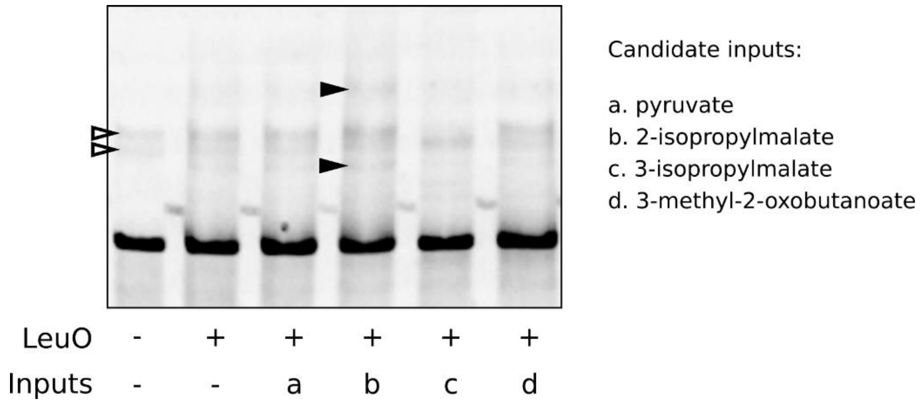


**Figure S4: Gel shift assay to identify LeuO signal molecule.** A representative gel is shown out of four independent gel shift experiments with 0 (-) or 25 nM (+) of LeuO protein and 0.5 nM of DNA fragment containing the *leuLABCD* promoter region. 10mM candidate inputs were added (+) to test their effect on LeuO binding. Hollow arrows indicate faint bands with unspecific shifts independent of LeuO. Black arrows indicate specific band shifts dependent of LeuO addition.
